# Dual-Mode Bio-CM^2^: Multimodal Computational Miniature Mesoscope for Fluorescence and Label-Free Imaging

**DOI:** 10.64898/2026.09.08.750281

**Authors:** Qilin Deng, Guorong Hu, Bradley C. Rauscher, Marcelo Villafuerte, Bethany Weinberg, Zhixiong Chen, Hui Feng, Anna Devor, Martin Thunemann, Lei Tian

## Abstract

Understanding biological function often requires complementary information on molecular identity, morphology, behavior, and physiology acquired simultaneously from the same specimen. Achieving this capability in miniature microscopes remains challenging because integrating multiple imaging contrasts typically increases optical complexity while compromising field of view (FOV), spatial resolution, or form factor. Here we present dual-mode Bio-CM^2^, a multimodal computational miniature mesoscope that simultaneously captures fluorescence and label-free reflectance through a shared optical architecture based on distributed computational optics. The system enables frame-interleaved, co-registered dual-mode imaging over a *∼* 7.5 *×* 10 mm^2^ FOV with *∼* 6 *µ*m lateral resolution. We demonstrate the versatility of dual-mode Bio-CM^2^ across diverse biological systems. In freely moving *Caenorhabditis elegans*, reflectance captures body posture and locomotor behavior that contextualize fluorescently labeled protein aggregates. In freely swimming larval zebrafish, reflectance captures whole-body morphology and swimming behavior, while fluorescence visualizes cardiac activity. In head-fixed mice, simultaneous fluorescence and reflectance imaging across the bilateral dorsal cortex provides complementary measurements of neuronal calcium activity and intrinsic hemodynamic signals. By integrating molecularly specific fluorescence with complementary reflectance contrast, dual-mode Bio-CM^2^ extends distributed computational optics into a scalable platform for multimodal miniature imaging.

## 1 Introduction

Biological processes span multiple spatial scales, from molecular signaling within individual cells to tissue organization, organ physiology, and animal behavior. Understanding these processes therefore requires complementary measurements that capture both molecular specificity and the surrounding structural, behavioral, and physiological context. Fluorescence microscopy provides molecular specificity through genetically encoded or exogenous labels [1], whereas label-free optical contrast reveals tissue morphology, vascular architecture, and intrinsic physiological dynamics [2]. Together, these complementary imaging contrasts provide a more complete view of biological function than either modality alone.

Translating these complementary measurements into miniature microscopes (miniscopes) suitable for freely behaving animals remains challenging because expanding imaging capability typically requires additional optical complexity while preserving a compact form factor [3, 4]. Recent advances in miniature optics and computational imaging have substantially expanded the field of view (FOV) and imaging performance of miniscopes [5–15]. We recently proposed *distributed computational optics*, a scalable framework that partitions optical image formation across multiple coordinated imaging modules and computationally integrates their measurements into a unified image, allowing optical complexity to scale with individual subfields rather than the entire FOV [16]. Bio-CM^2^ was the first realization of this framework, enabling cellular-resolution fluorescence imaging from individual cortical vessels to thousands of neurons across nearly the entire mouse dorsal cortex while overcoming the conventional trade-off between FOV and spatial resolution.

Having established a scalable architecture for large-FOV miniature fluorescence imaging, the next challenge is to expand the diversity of biological information captured within the same platform. While Bio-CM^2^ demonstrated distributed computational optics for fluorescence imaging, it retained the single-contrast paradigm shared by most existing miniscopes. Recent dual-color fluorescence miniscopes have expanded molecular multiplexing by simultaneously imaging multiple fluorescent reporters [8, 17, 18]. A complementary strategy is to combine fluorescence with label-free reflectance, enabling molecularly specific fluorescence to be interpreted alongside complementary structural and physiological information.

The complementary value of fluorescence and label-free reflectance has been demonstrated across diverse imaging modalities. In wide-field optical mapping, reflectance monitors cerebral hemodynamics and corrects fluorescence signals for hemoglobin absorption artifacts [19–21]. This concept was subsequently translated to miniature microscopy through mini-mScope, enabling simultaneous fluorescence and reflectance imaging of the dorsal cortex in freely behaving mice [22]. Dual-mode fluorescence– reflectance imaging has also been explored in endomicroscopy for *in situ* cancer diagnosis, combining molecular and label-free tissue contrast [23]. These studies highlight the biological value of complementary fluorescence and reflectance imaging, but integrating both modalities while preserving large FOV, cellular resolution, and a compact miniature form factor remains challenging.

Here, we present dual-mode Bio-CM^2^, which extends the distributed computational optics framework to integrate fluorescence and label-free reflectance within a shared optical architecture. The resulting system preserves the large FOV and cellular-resolution imaging performance of Bio-CM^2^ while enabling co-registered dual-mode imaging in a compact miniature mesoscope. We demonstrate the versatility of dual-mode Bio-CM^2^ across diverse biological systems. In freely moving *Caenorhabditis elegans*, fluorescence visualizes protein aggregates while reflectance captures body posture and locomotor behavior. In freely swimming larval zebrafish, fluorescence localizes cardiac tissue while reflectance captures whole-body morphology and swimming trajectories. In mouse cortex, fluorescence reports neuronal calcium activity while reflectance measures cortical hemodynamics. Together, these demonstrations establish distributed computational optics as a scalable framework for multimodal miniature imaging.

## 2 Results

### 2.1 Design of Dual-Mode Bio-CM^2^

We developed dual-mode Bio-CM^2^, a miniature mesoscope that integrates fluorescence and label-free reflectance imaging within a shared optical architecture (Figure 1A). Both imaging modalities share the same collection optics, emission filter, and image sensor, enabling molecularly specific fluorescence and label-free structural information to be acquired from the same FOV without beam splitters, duplicated detection paths, or post-acquisition image registration. This shared architecture preserves a compact form factor while intrinsically co-registering the two imaging modalities.

**Figure 1.**
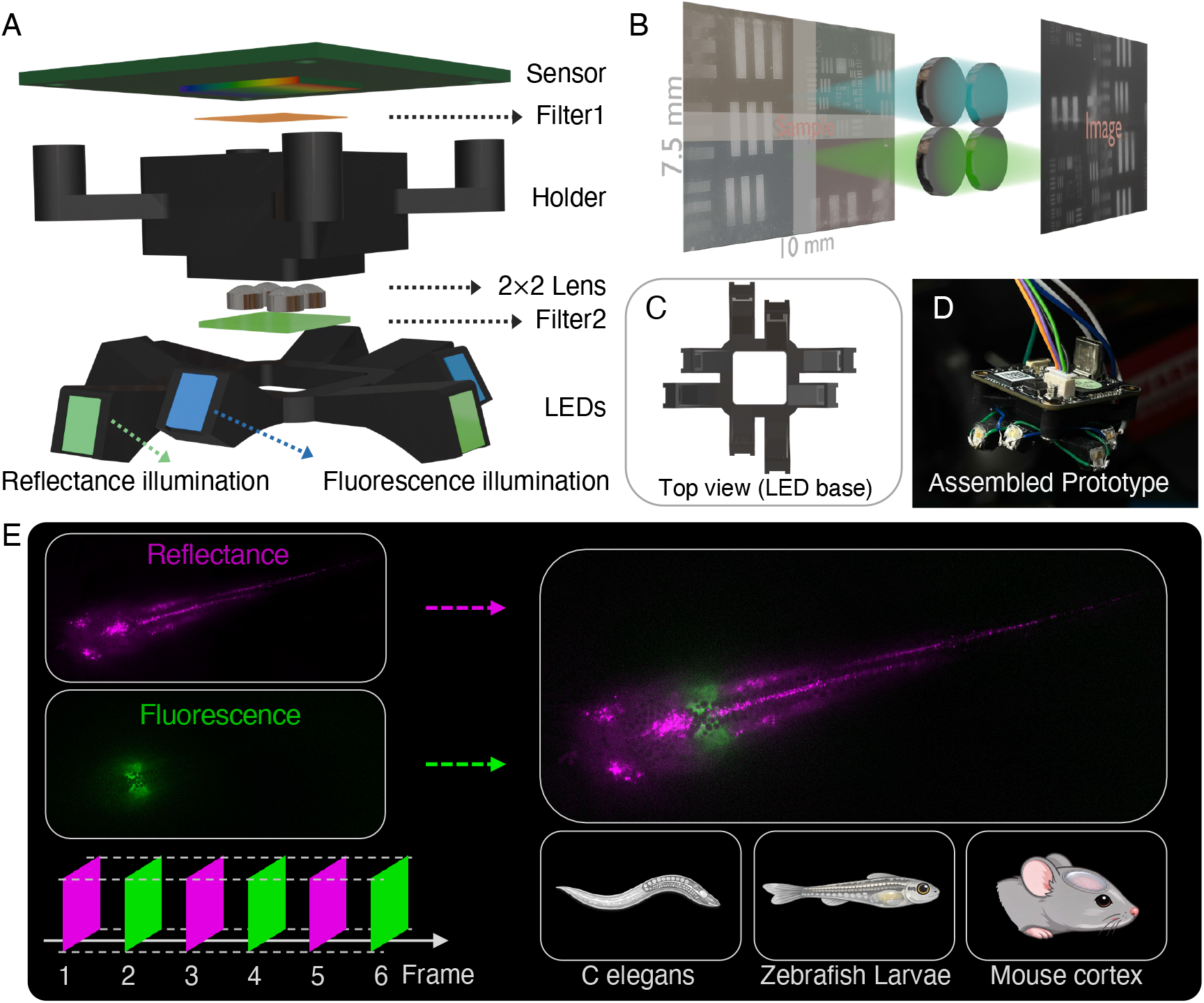
Overview of dual-mode Bio-CM^2^. Dual-mode Bio-CM^2^ extends the distributed computational optics architecture of Bio-CM^2^ to enable simultaneous fluorescence and label-free reflectance imaging within a compact miniature microscope. (A–D) System architecture. (A) Exploded view of the miniature imaging head. (B) Distributed computational optics architecture, in which four miniature objectives image overlapping subregions onto a common image sensor that are computationally stitched into an approximately 7.5 *×* 10 mm^2^ field of view. (C) Top view of the dual-wavelength illumination module surrounding the shared imaging path. (D) Photograph of the assembled prototype. (E) Operating principle and biological applications. Frame-interleaved acquisition alternates fluorescence and reflectance imaging through the shared optical path to produce intrinsically co-registered multimodal images across diverse biological systems, including *C. elegans*, zebrafish larvae, and the mouse brain.

The large FOV is enabled by the distributed computational optics architecture established in Bio-CM^2^ [16], in which four miniature imaging modules simultaneously acquire overlapping subregions that are computationally stitched into a continuous image (Figure 1B). This architecture maintains cellular-scale resolution over an approximately 7.5 *×* 10 mm^2^ FOV while preserving the compactness required for miniature imaging. Here, we extend this platform to complementary fluorescence and label-free reflectance imaging.

Label-free reflectance imaging is enabled through the co-design of the illumination geometry and spectral configuration. The reflectance illumination is delivered at a steep off-axis angle that directs specular reflections from planar samples outside the collection aperture while efficiently collecting scattered light across the full FOV. The reflectance wavelength is selected within the fluorescence emission passband, allowing reflected light and fluorescence emission to share the same detection path while remaining spectrally separated at excitation.

Fluorescence and reflectance images are acquired using frame-interleaved illumination synchronized to the camera timing (Figure 1E). Alternating excitation wavelengths assign one imaging modality to each camera frame, while temporally gated illumination produces a pseudo-global-shutter acquisition that minimizes rolling-shutter distortion during imaging of dynamic specimens. Together, these design elements enable near-simultaneous fluorescence and label-free imaging over a large FOV while providing intrinsically co-registered molecular and structural information.

### 2.2 Optical characterization

We characterized the optical performance of dual-mode Bio-CM^2^ through measurements of FOV, spatial resolution, and spectral response (Figure 2). Imaging a USAF 1951 resolution target yielded an FOV of approximately 7.5 *×* 10 mm^2^, substantially larger than that of a conventional 0.1 NA, 2*×* tabletop microscope shown at the same scale for comparison (Figure 2A). This expanded FOV enables centimeter-scale specimens to be imaged without mechanical scanning.

**Figure 2.**
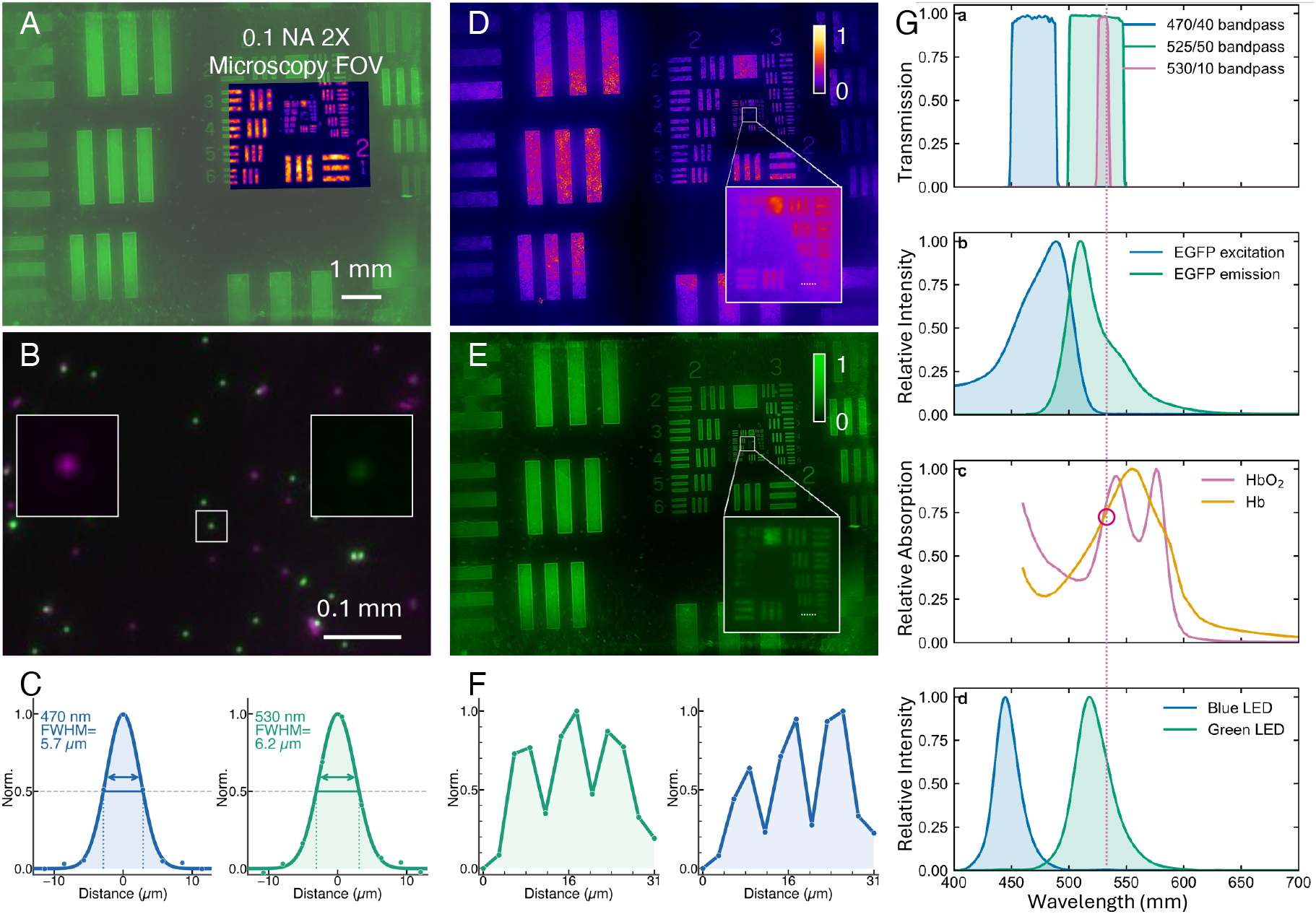
Optical and spectral characterization of dual-mode Bio-CM^2^. (A) USAF 1951 resolution target imaged over the approximately 7.5 *×* 10 mm^2^ FOV of dual-mode Bio-CM^2^ under 530 nm reflectance illumination. The inset shows the FOV of a conventional 0.1 NA, 2*×* microscope objective at the same physical scale. (B) Dual-channel image of a mixed bead sample containing 4 *µ*m green- and red-fluorescent beads. Insets show magnified views of a single green-fluorescent bead in the fluorescence and reflectance channels. (C) Lateral resolution measured using 4 *µ*m green-fluorescent beads, yielding FWHM values of 5.7 *µ*m in fluorescence and 6.2 *µ*m in reflectance. (D,E) Fluorescence (D) and reflectance (E) images of a fluorescent USAF 1951 resolution target. Insets show magnified views of the boxed Group 6 regions. (F) Intensity profiles across the Group 6, Element 6 bar pattern indicated in (D) and (E), resolving the three 4.4 *µ*m-wide bars in both imaging modalities. (G) Spectral configuration of dual-mode Bio-CM^2^ showing (a) transmission spectra of the excitation, emission, and reflectance filters, (b) EGFP excitation and emission spectra, (c) absorption spectra of oxyhemoglobin (HbO2) and deoxyhemoglobin (Hb), and (d) measured spectra of the blue and green LEDs. Vertical dashed lines indicate the 530 nm reflectance wavelength.

To illustrate the complementary image contrast provided by the two modalities, we imaged a mixed bead sample containing 4 *µ*m green- and red-fluorescent beads (Figure 2B). Green fluorescent beads appear white in the merged image because they contribute to both the fluorescence (green) and reflectance (magenta) channels. In contrast, red fluorescent beads appear only in the reflectance channel because their fluorescence emission lies outside the detection band. This experiment illustrates the complementary contrast of the two imaging modalities.

We next quantified the spatial resolution of each imaging modality using 4 *µ*m green-fluorescent beads imaged sequentially in fluorescence and reflectance modes. Gaussian fits to individual bead intensity profiles yielded lateral full-width-at-half-maximum (FWHM) values of 5.7 *µ*m for fluorescence and 6.2 *µ*m for reflectance (Figure 2C). Resolution was further validated using a fluorescent USAF 1951 target (Figure 2D–F). Both imaging modalities resolved Group 6, Element 6, corresponding to a 4.4 *µ*m line width, with line profiles showing three clearly separated peaks. Resolution beyond Group 6, Element 6 was limited by reduced target contrast and residual off-axis aberrations rather than the intrinsic optical resolution. The measured performance is consistent with the sampling limit set by the effective system magnification and sensor pixel size. Spatial variations in the point spread function (PSF) across the FOV are further characterized in *SI Appendix*, Fig. S2.

Finally, we characterized the spectral configuration underlying dual-mode imaging (Figure 2G). The reflectance wavelength was selected to satisfy both optical and biological design considerations. Optically, the 530/10 bandpass filter confines the reflectance illumination to a narrow spectral band within the transmission band of the shared ET525/50m emission filter (Figure 2G(a,d)). Consequently, EGFP fluorescence and reflected green light share the same detection path while remaining spectrally separated from excitation (Figure 2G(b)).

Biologically, 530 nm lies near an isosbestic point of oxyhemoglobin (HbO_2_) and deoxyhemoglobin (Hb), where both chromophores exhibit similar absorption coefficients (Figure 2G(c)) [24]. Consequently, diffuse reflectance at this wavelength is primarily sensitive to changes in total hemoglobin (HbT), providing a label-free measure of cerebral blood volume [19, 25]. We therefore selected 530 nm to complement GCaMP fluorescence imaging with label-free hemodynamic contrast while preserving a shared optical detection path. Although 530 nm was selected here for optical compatibility and hemodynamic imaging, the dual-mode Bio-CM^2^ architecture can readily accommodate other reflectance wavelengths to target alternative endogenous contrast mechanisms.

### 2.3 Dual-mode acquisition characterization

We validated that frame-interleaved acquisition effectively separates fluorescence and reflectance signals while preserving spatial co-registration (Figure 3). Because the 530 nm reflectance illumination shares the fluorescence detection band, suppression of specular reflection relies on off-axis dark-field illumination rather than spectral filtering. The off-axis reflectance illumination directs specular reflections outside the collection aperture, producing a dark-field imaging configuration, whereas fluorescence excitation uses epi-illumination, with reflected excitation light rejected by the emission filter (Figure 3A). Imaging a polished mirror confirmed this behavior, revealing only surface scratches and dust without detectable specular background (*SI Appendix*, Fig. S3).

**Figure 3.**
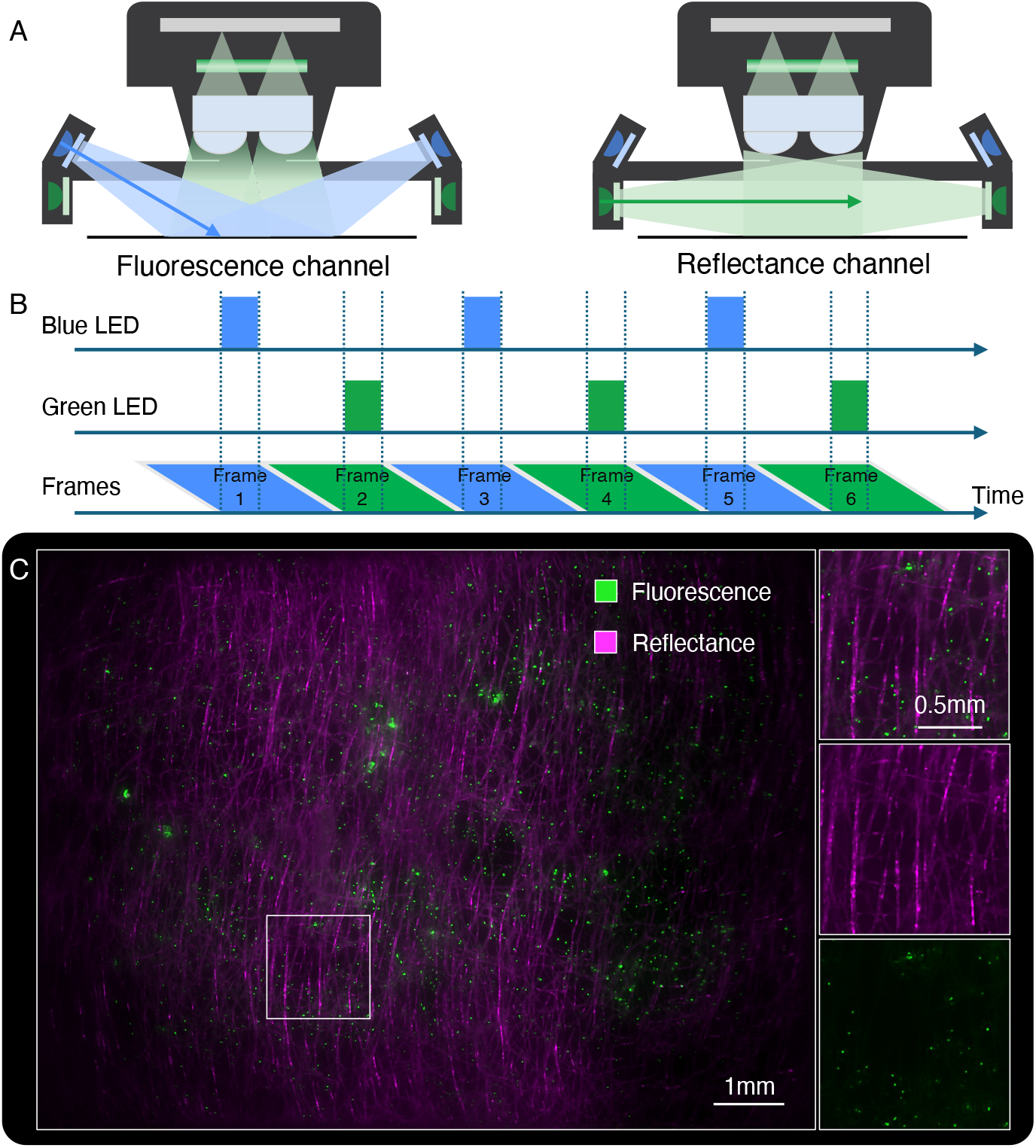
Dual-mode acquisition in Bio-CM^2^. (A) Illumination geometries for fluorescence (left) and reflectance (right) imaging. Fluorescence excitation is rejected by the emission filter, whereas reflectance illumination is delivered at a steeper angle to suppress specular reflections. (B) Frame-interleaved acquisition synchronized with the rolling-shutter CMOS sensor. Blue and green LED pulses alternate between successive camera frames. The slanted parallelograms represent rolling-shutter exposure, with successive sensor rows integrating sequentially. The shortened LED pulses are confined to the common integration interval, ensuring that each frame contains only a single illumination wavelength. (C) Dual-channel imaging of a bead–fiber phantom containing 4 *µ*m green-fluorescent beads embedded in a non-fluorescent paper fiber network. The boxed region is enlarged to show the merged image (top), reflectance channel (middle), and fluorescence channel (bottom).

We then evaluated temporal separation of the two imaging modes (Figure 3B). Because the CMOS sensor employs a rolling shutter, blue and green LED pulses were synchronized to the interval during which all sensor rows integrate simultaneously and switched off during sensor readout. This timing ensures that each frame contains only one illumination wavelength, minimizing channel cross-talk while producing a 20 frames-per-second (fps) acquisition stream with 10 fps per imaging modality (*SI Appendix*, Section 4).

Finally, we validated dual-mode imaging using a bead–fiber phantom containing fluorescent beads embedded in a non-fluorescent scattering fiber network (Figure 3C). The merged image combines fluorescence and reflectance information from the same FOV, while the individual channels recover complementary contrast: the fluorescence channel selectively visualizes the fluorescent beads, whereas the reflectance channel resolves the fiber network. No detectable cross-talk was observed between the two channels, validating accurate channel separation together with intrinsic spatial co-registration.

### 2.4 Dual-mode imaging of freely swimming *C. elegans*

We first evaluated whether dual-mode imaging provides complementary molecular and behavioral information in freely moving animals using transgenic *Caenorhabditis elegans* strain AM141 [*unc-54p::Q40::YFP*], which expresses 40 polyglutamine repeats fused to yellow fluorescent protein (YFP) in body-wall muscle as a Huntington’s-disease-associated aggregation model (Figure 4 and Movie S1) [26]. *C. elegans* is a widely used model for studying the relationship between molecular pathology and behavior, where imaging during natural locomotion is often essential. Because fluorescence labels only the molecular target of interest, interpreting these signals requires simultaneous knowledge of the animal’s body configuration. Dual-mode imaging provides both measurements from the same acquisition.

**Figure 4.**
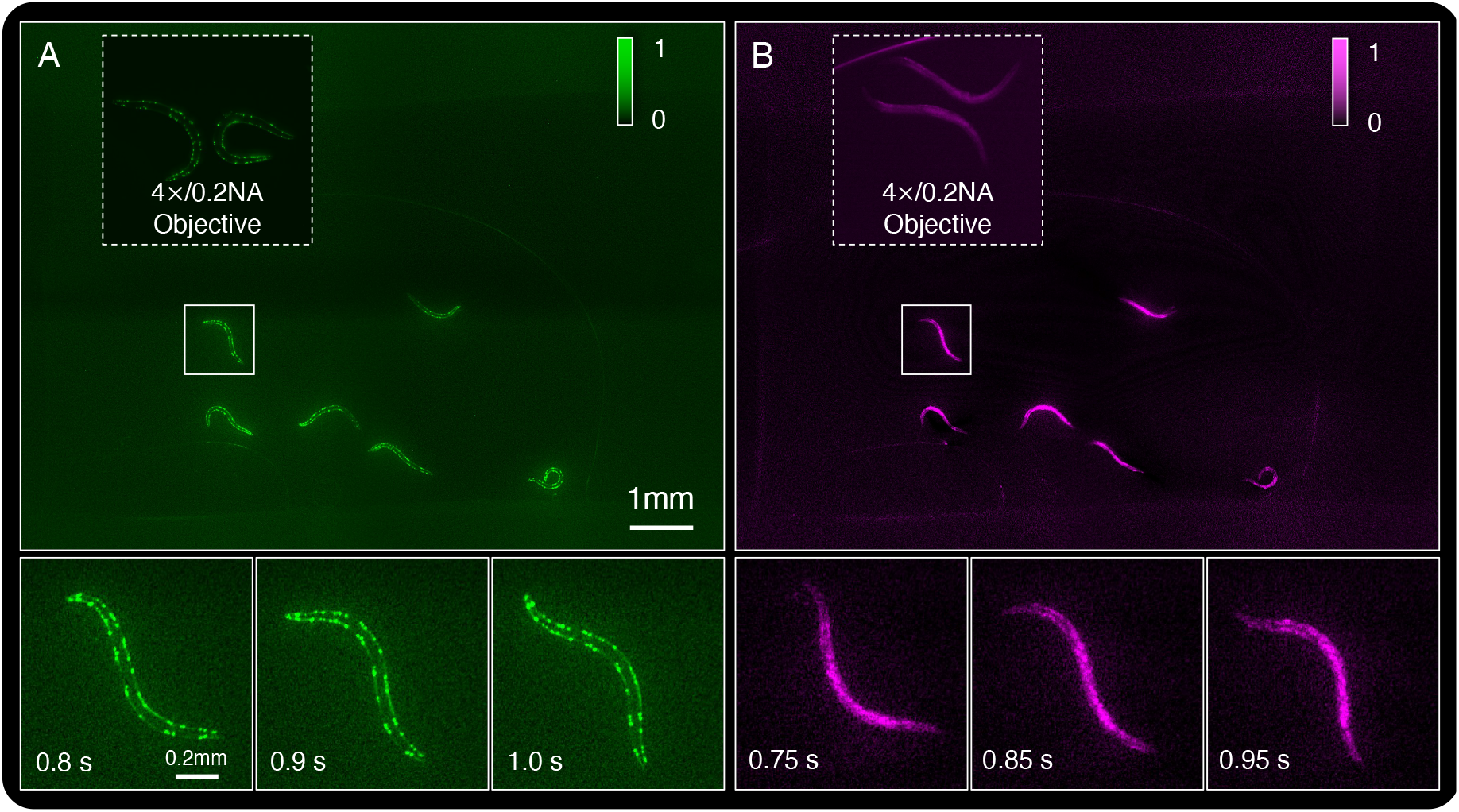
Dual-mode imaging of freely swimming YFP-labeled *C. elegans*. (A) Fluorescence-channel image of multiple freely swimming *C. elegans* strain AM141[*unc-54p::Q40::YFP*]. The inset (top left) shows the same specimen imaged with a Nikon Plan Apo *λ* 4*×*/0.2 NA objective at the same physical scale. The dashed box indicates the worm shown in the time-lapse sequence below, with representative fluorescence frames at *t* = 0.8, 0.9, and 1.0 s. (B) Frame-interleaved reflectance-channel image of the same field. The inset shows the corresponding benchtop reflectance image acquired with the same objective. The dashed box marks the same worm as in (A). Representative reflectance frames at *t* = 0.75, 0.85, and 0.95 s are shown below. See also Movie S1.

We imaged multiple freely swimming worms within a single FOV. The fluorescence channel (Figure 4A) resolved YFP-labeled polyglutamine aggregates distributed along the body-wall muscle, whereas the simultaneously acquired reflectance channel (Figure 4B) delineated the complete body contour and bending posture. Since the AM141 fluorescence consists primarily of discrete aggregate puncta rather than continuous body labeling, the reflectance image provides the anatomical framework needed to localize these aggregates throughout locomotion. The benchtop comparison inset acquired with a Nikon Plan Apo *λ* 4*×*/0.2 NA objective illustrates the substantially smaller imaging area of a conventional microscope despite comparable spatial resolution.

Frame-interleaved acquisition recorded both channels at 10 fps with alternating fluorescence and reflectance exposures separated by 50 ms (Figure 4, lower panels). Fluorescence frames at *t* = 0.8, 0.9, and 1.0 s captured the YFP-labeled aggregates as a representative worm executed a body-bending stroke, while the interleaved reflectance frames (*t* = 0.75, 0.85, and 0.95 s) resolved the corresponding body posture throughout the same locomotor cycle. The alternating measurements provide temporally registered molecular and morphological information throughout the locomotor cycle.

Together, these results demonstrate that dual-mode Bio-CM^2^ simultaneously captures molecularly specific fluorescence and label-free morphology from freely behaving organisms. The complementary measurements place sparse molecular signals within their anatomical and behavioral context, enabling direct interpretation of molecular phenotypes during natural locomotion.

### 2.5 Dual-mode imaging of freely swimming zebrafish

We next evaluated dual-mode imaging in a freely swimming vertebrate using a transgenic larval zebrafish (approximately 12 d post-fertilization, 5.34 mm body length) from the compound line *Tg(dβh:MYCN; dβh:EGFP; cmlc2:EGFP)*, which expresses EGFP in the heart and neural crest cells (Figure 5 and Movie S2). Larval zebrafish are a widely used model for linking organ-specific physiology to whole-animal behavior. Because fluorescence labels only genetically defined tissues, interpreting these signals during free swimming requires simultaneous visualization of the animal’s morphology and locomotion.

**Figure 5.**
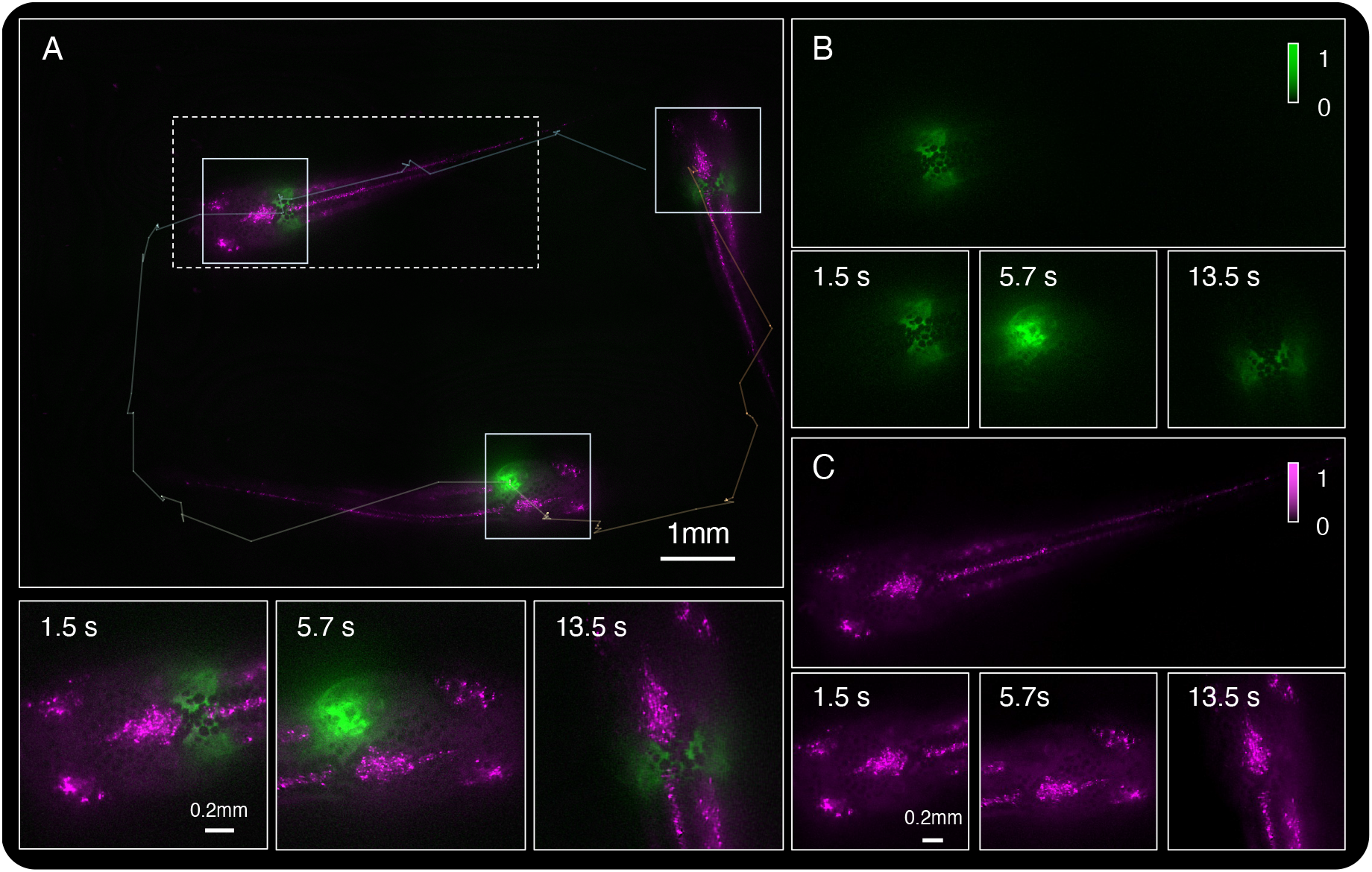
Dual-mode imaging of a freely swimming transgenic larval zebrafish. (A) Merged fluorescence–reflectance image showing the zebrafish at three representative time points during free swimming (*t* = 1.5, 5.7, and 13.5 s). Fluorescence is rendered in green and reflectance in magenta. The tracked swimming trajectory is overlaid in white. Solid white boxes mark the fish at the corresponding time points, and the dashed box outlines the region enlarged in the lower row. Bottom: enlarged merged images at the corresponding time points. (B) Representative fluorescence image showing EGFP expression predominantly in the heart (green, intensity normalized to 0–1). Bottom: fluorescence images at *t* = 1.5, 5.7, and 13.5 s. (C) Representative reflectance image revealing whole-body morphology through dark-field scattering (magenta, intensity normalized to 0–1). Bottom: reflectance images at *t* = 1.5, 5.7, and 13.5 s, showing body posture during locomotion. See also Movie S2 (real-time playback, 10 fps per channel).

Fish were imaged in a custom 3D-printed arena (8 *×* 11 *×* 2 mm) that matched the imaging FOV while maintaining the animal within the system’s depth of focus. The fluorescence channel (Figure 5B) selectively visualized EGFP expression in the heart, with weaker expression in neural crest cells, whereas the frame-interleaved reflectance channel (Figure 5C) continuously captured the animal’s whole-body morphology, including the head, trunk, tail, body posture, and swimming trajectory. The merged dual-mode images localized the fluorescence signal within the structural context of the freely moving animal, allowing organ-specific fluorescence to be interpreted together with whole-body morphology and behavior.

Representative images acquired at three time points (*t* = 1.5, 5.7, and 13.5 s) illustrate continuous dual-mode imaging throughout free swimming (Figure 5A–C). The overlaid merged images visualize the fish at successive positions along its swimming path, while the corresponding fluorescence and reflectance images separately highlight localized EGFP expression and whole-body morphology. Enlarged views further demonstrate accurate co-registration of the two imaging modalities throughout locomotion.

Together, these results demonstrate that dual-mode Bio-CM^2^ enables simultaneous imaging of organ-specific fluorescence and whole-body behavior in freely swimming vertebrates. By combining localized cardiac fluorescence with continuous measurements of body posture and swimming trajectories over the entire behavioral arena, the platform links tissue-specific molecular signals to animal-scale behavior within a single miniature imaging system.

### 2.6 Dual-mode imaging of neuronal and hemodynamic activity across the mouse cortex

Finally, we leveraged the large FOV of dual-mode Bio-CM^2^ to perform fluorescence–reflectance imaging of neuronal and hemodynamic signals across nearly the entire mouse dorsal cortex within a single recording (Figure 6A,B; Movie S3). We imaged a head-fixed mouse expressing GCaMP6s in layer 5 pyramidal neurons (Rbp4-Cre injected with AAV-PHP.eB.GCaMP6s) through a glass cranial window (Figure 6D). Interleaved 470 nm fluorescence excitation and 530 nm reflectance illumination produced two intrinsically co-registered image streams: GCaMP fluorescence (Δ*F/F*) (Figure 6A), reporting neuronal calcium activity, and reflectance (Δ*R/R*) (Figure 6B), providing a label-free hemodynamic readout sensitive to blood-volume changes near the hemoglobin isosbestic wavelength. The two modalities exhibit matching vascular anatomy, with major cortical vessels appearing as dark absorption features in both images, while localized functional signal changes display distinct spatial distributions, reflecting the complementary neuronal and hemodynamic contrast provided by the fluorescence and reflectance channels, respectively. Together, these complementary measurements enable simultaneous mapping of neuronal calcium activity and instrinsic hemodynamic signals across the cortex while providing the physiological reference required for subsequent reflectance-based correction of hemodynamic contamination in the fluorescence signal [19, 25].

**Figure 6.**
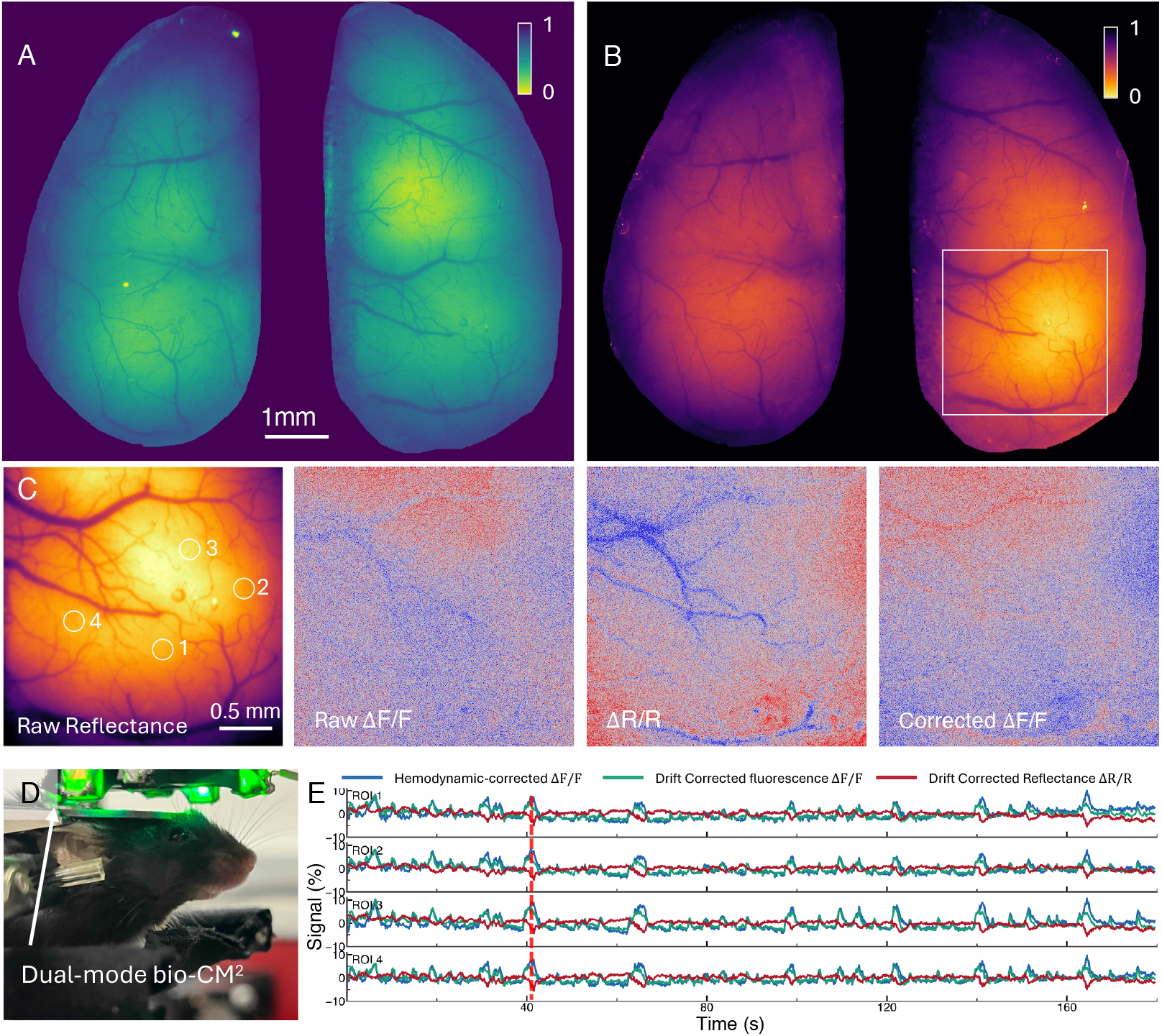
Dual-mode wide-field optical mapping of bilateral mouse cortex with hemodynamic correction. (A) Single-frame GCaMP6s fluorescence image of the bilateral cortex acquired in a head-fixed mouse expressing GCaMP6s in layer 5 pyramidal neurons. Cortical vasculature appears dark because hemoglobin attenuates both excitation and emission light. (B) Simultaneously acquired green reflectance image of the same FOV, providing intrinsic hemodynamic contrast. The white box indicates the enlarged region shown in C, and white circles mark the four ROIs analyzed in E. (C) Hemodynamic correction of the boxed cortical region at *t* = 41.6 s. From left to right: raw reflectance image; raw fluorescence Δ*F/F* ; reflectance Δ*R/R*; and hemodynamic-corrected fluorescence obtained by pixelwise ratiometric correction (Eq. (6)). The vascular pattern is substantially reduced after correction. (D) Photograph of the head-fixed mouse with a bilateral glass cranial window. (E) Frame-interleaved fluorescence and reflectance imaging with corresponding traces from the four ROIs during spontaneous resting-state activity (170 s, 10 fps per channel). Drift-corrected fluorescence (Δ*F/F*) and reflectance (Δ*R/R*) traces are shown together with the hemodynamic-corrected fluorescence trace. The dashed line indicates the time point shown in (C). Fluorescence and reflectance co-vary because of shared hemodynamic fluctuations, whereas ratiometric correction suppresses this component while preserving neural transients. See also Movie S3 for the frame-interleaved recording.

A central challenge in estimating neuronal activity from cortex-wide GCaMP imaging is that hemoglobin absorption attenuates fluorescence along both the excitation and emission light paths [19, 25]. Consequently, changes in blood volume arising from either systemic physiology or neurovascular coupling modulate the measured fluorescence signal and can be mistaken for changes in neuronal activity [27]. Wide-field optical mapping therefore commonly acquires a hemodynamic-sensitive reflectance signal along with fluorescence to correct for this vascular contamination.

Figure 6C demonstrates the spatial effect of hemodynamic correction at a representative time point (*t* = 41.6 s). The raw GCaMP Δ*F/F* map (Figure 6C, second) exhibits broad vascular-associated intensity variations that spatially correspond to the vasculature visible in the reflectance image (Figure 6C, first) and the hemodynamic Δ*R/R* map (Figure 6C, third). Applying pixel-wise ratiometric correction substantially suppresses these vascular-associated fluctuations, producing a corrected Δ*F/F* map (Figure 6C, fourth) in which neuronal activity-related contrast is more clearly resolved.

The temporal effect of the correction is illustrated by four representative cortical ROIs (Figure 6E). During *∼* 170 s of spontaneous resting-state activity, the raw GCaMP Δ*F/F* and reflectance Δ*R/R* traces exhibited correlated slow fluctuations characteristic of shared hemodynamic modulation. Both raw traces also showed a gradual downward drift over time (Supplementary Fig. 7), consistent with reduced illumination intensity caused by progressive LED heating during the recording. Drift correction removed this slow channel-wide intensity trend while preserving faster fluorescence and reflectance dynamics. A subsequent reflectance-based ratiometric correction attenuated fluorescence fluctuations that covaried with the 530 nm reflectance signal, thereby reducing vascular contamination while preserving transient neuronal calcium events.

In a complementary recording with periodic air-puff whisker stimulation, the temporal relationship between neuronal and hemodynamic signals was further quantified using cross-correlation and magnitude-squared coherence analyses (*SI Appendix*, Fig. S6 and Movie S4). The reflectance signal consistently followed the GCaMP fluorescence signal by approximately 1.1 s, consistent with the expected delay of the hemodynamic response relative to neuronal activity [25]. Furthermore, the two signals exhibited near-unity coherence (0.97–0.98) within the slow (*<* 0.1 Hz) hemodynamic band, demonstrating strong neurovascular coupling and confirming that the simultaneously acquired reflectance channel faithfully captures physiologically relevant hemodynamic signals.

Together, these results demonstrate that dual-mode Bio-CM^2^ extends distributed computational optics to multimodal functional imaging, translating the complementary capabilities of wide-field optical mapping into a compact miniature platform.

## 3 Discussion

We developed dual-mode Bio-CM^2^ by extending the Bio-CM^2^ platform from single-contrast fluorescence imaging to multimodal fluorescence–reflectance imaging. Rather than increasing molecular multiplexing through additional fluorescent labels, the system combines fluorescence with complementary label-free reflectance to simultaneously capture molecular specificity together with structural, behavioral, and physiological context. Across freely swimming *C. elegans*, larval zebrafish, and mouse cortex, the reflectance channel contributed information that fluorescence alone could not provide, demonstrating the versatility of complementary multimodal imaging across diverse biological systems.

The present work extends Bio-CM^2^ along a different dimension from the original platform. Bio-CM^2^ addressed the spatial trade-off between FOV and resolution through distributed computational optics, enabling cellular-resolution fluorescence imaging across nearly the entire mouse dorsal cortex. Here, we address a complementary challenge—expanding the biological information content of each measurement. By integrating fluorescence and dark-field reflectance within the same distributed computational imaging architecture, the platform preserves the large FOV and cellular-scale imaging performance of Bio-CM^2^ while providing co-registered molecular and label-free measurements. More broadly, these results demonstrate that distributed computational optics provides a flexible platform for incorporating complementary imaging modalities without fundamentally changing the underlying imaging architecture.

Fluorescence and reflectance are acquired sequentially by frame interleaving rather than simultaneously. Although this acquisition scheme was sufficient for the biological demonstrations presented here, applications requiring precise synchronization of rapid dynamics may benefit from simultaneous dual-channel acquisition. The current prototype was designed as a proof of concept rather than a fully optimized head-mounted device, and the cortical imaging experiments were therefore performed under head-fixed conditions. Future work will focus on further miniaturization and weight reduction while expanding the multimodal imaging capability of the platform.

The modular architecture of Bio-CM^2^ naturally supports future extensions beyond the present implementation. Additional fluorescence channels could increase molecular multiplexing, while multi-wavelength reflectance could enable quantitative measurements of cerebral blood volume and oxygenation [19, 25]. More broadly, the ability to integrate complementary imaging modalities within a shared distributed computational optics architecture provides a foundation for compact miniature microscopes that simultaneously capture multiple aspects of biological function while preserving large FOV and cellular-scale resolution.

## Methods

### Dual-mode Bio-CM^2^ system design and implementation

Dual-mode Bio-CM^2^ is built around a monochrome CMOS camera (DMM 37UX226-ML, The Imaging Source) with a 1.85 *µ*m pixel size. The imaging optics consist of a 2 *×* 2 array of four identical miniature plano-aspheric lenses (#16-684, Edmund Optics), each imaging an overlapping subregion of the sample onto one quadrant of the shared image sensor (Figure 1B). The lens array provides an effective numerical aperture (NA) of approximately 0.17 and a system magnification of approximately 0.65*×*, corresponding to a sample-plane pixel size of approximately 2.85 *µ*m. The overlapping measurements from the four imaging modules are computationally stitched to form a continuous large-field image.

A single fluorescence emission filter (ET525/50m, Chroma; 500–550 nm passband) is shared by both fluorescence and reflectance imaging channels and mounted between the lens array and the image sensor, eliminating the need for a dichroic beam splitter or a second camera. Because interference filters exhibit angle-dependent spectral shifts at oblique incidence, an additional long-pass absorption filter (#54-467, Edmund Optics; cutoff wavelength *∼*500 nm) is mounted in series with the emission filter to suppress angle-dependent spectral leakage.

Illumination is provided by eight LEDs arranged symmetrically around the imaging path (Figure 1A,C): four blue LEDs (LXML-PB01-0040, Lumileds) for fluorescence excitation and four green LEDs (LXML-PM01-0100, Lumileds) for label-free reflectance, with one blue and one green LED mounted on each side of the square. The blue LEDs are filtered by 470/40 nm excitation bandpass filters (ET470/40x, Chroma) to excite GFP-, YFP-, and GCaMP-based fluorophores. To enable fluorescence and reflectance imaging to share the same collection optics and image sensor, the green LEDs are filtered by a 530/10 nm bandpass filter (Chroma), so that the reflectance illumination lies within the passband of the shared fluorescence emission filter. Blue illumination is delivered in an epi-illumination geometry, whereas green illumination is introduced at a steep off-axis angle such that specular reflections from a planar sample are directed outside the collection aperture (Figure 3A).

The optomechanical housing, lens-array holder, and LED mount are designed and fabricated by stereolithographic 3D printing (Black Resin V4.1, Formlabs).

An Arduino Uno R3 monitors the camera strobe output and alternately switches the blue and green LED arrays on successive camera frames, producing frame-interleaved fluorescence and reflectance acquisition at 10 frames s*^−^*^1^ per channel (20 frames s*^−^*^1^ total). To suppress rolling-shutter distortion, a brief LED pulse is emitted during the interval in which all sensor rows are simultaneously integrating. Because every row records light from the same instant in time, this illumination-gated acquisition produces a pseudo-global shutter while maintaining frame-interleaved dual-mode imaging. Details of the synchronization electronics and rolling-shutter timing are provided in *SI Appendix*, Figs. S4 and S5. A complete bill of materials is provided in *SI Appendix*, Table S1.

### Sub-image registration and stitching

Each camera frame was partitioned into four sub-images corresponding to the four imaging modules in the 2*×*2 lens array. Each sub-image underwent background correction, geometric distortion correction, and field-dependent deconvolution using experimentally measured point spread functions (PSFs). The restored sub-images were then stitched into a single composite image.

Image stitching was based on a one-time geometric calibration using a USAF 1951 resolution target spanning the full FOV. Spatial transformations for each imaging module were estimated from corresponding features in the overlapping regions and subsequently applied to all recordings without per-frame optimization. The overlap between adjacent sub-images enabled accurate registration and seamless blending across module boundaries. A schematic of the reconstruction pipeline is shown in *SI Appendix*, Fig. S1. Additional implementation details are described in [16].

### *C. elegans* strain and culture

Transgenic *Caenorhabditis elegans* AM141[*unc-54p::Q40::YFP*], expressing 40 polyglutamine repeats fused to yellow fluorescent protein under the body-wall-muscle *unc-54* promoter [26], was used for the worm imaging experiments and was obtained from the *Caenorhabditis* Genetics Center (CGC) without further modification. Worms were cultured at 15*^◦^*C on nematode growth medium (NGM) plates (per 1 L, 20 mM phosphate buffer, 0.8 mM CaCl_2_, 0.8 mM MgSO_4_, 17.5 g agar, 3.0 g NaCl, 2.5 g peptone, and 0.005 g cholesterol). Plates were seeded with OP50 *Escherichia coli* as a food source. Young adult worms from an unsynchronized population, day 1–3 of adulthood, were picked for imaging, and were fed and healthy at the time of imaging. For mounting, double-sided tape was adhered to a microscope slide, and a razor was used to create a 1.5 *×* 0.5 cm channel in the tape. Worms were transferred into *∼*10 *µ*L of water within the channel and sealed under a coverslip over the double-sided tape for free-swimming recordings with the dual-mode miniscope.

### Zebrafish husbandry

Zebrafish husbandry and handling were performed in the aquatic facility at Boston University Chobanian & Avedisian School of Medicine under approved protocols from the Institutional Animal Care and Use Committee (IACUC). The Tg(cmlc2:EGFP) fish in the AB wild-type background were crossed to generate the compound transgenic Tg(cmlc2:EGFP) progeny used for imaging. In this line, EGFP served as the fluorescent indicator for the miniscope’s fluorescence channel, expressed under the *dβh* promoter in neural crest-derived cells and under the *cmlc2* promoter in the heart, while co-expression of MYCN under the same *dβh* promoter modeled MYCN-driven neuroblastoma. Larvae at approximately 12 d post-fertilization were used for free-swimming dual-mode imaging.

### Mice Preparation

All experimental procedures involving mice were performed in accordance with the guidelines established by the Boston University Institutional Animal Care and Use Committee. Mice were housed under a 12-h light/dark cycle, with lights on at 7:30 a.m., and were provided standard rodent chow and water *ad libitum*.

One adult female mouse from the Rbp4-Cre KL100Gsat line (MGI: 4367067) was used. Cortex-wide expression of GCaMP6s in layer 5 pyramidal neurons was induced by retro-orbital injection of an adeno-associated virus with the PHP.eB serotype [28]. The viral preparation was obtained from Addgene (#104495-PHPeB; pGP-AAV-CAG-FLEX-jGCaMP7s-WPRE) at a concentration of 3.2 *×* 10^13^ viral copies mL*^−^*^1^. A volume of 15 *µ*L was administered when the mouse was 6–8 weeks old.

The surgical procedure was adapted from previously described methods [29, 30]. Dexamethasone was administered approximately 4 h before surgery by intraperitoneal injection (4.8 mg kg*^−^*^1^ at a concentration of 4 mg mL*^−^*^1^) to minimize brain swelling associated with the surgical intervention. Slow-release buprenorphine and meloxicam were also administered before surgery. During surgery, the mouse was anesthetized with either isoflurane (2% in O_2_ for induction and 1% in O_2_ throughout the procedure), ketamine/xylazine, or a mixture of medetomidine, midazolam, and fentanyl. Body temperature was maintained at 37*^◦^*C throughout the procedure.

A custom-designed titanium headpost was attached to the cranium. The dorsal cranium was replaced with two pieces of curved or flat glass, one over each hemisphere. To reduce heat loss through the cranial window, either silicone or cotton was placed over the glass and enclosed by a protective three-dimensionally printed cap secured to the headpost. The mouse was allowed to recover for at least one week after surgery before habituation to head fixation and for at least three weeks before imaging experiments began.

Beginning at least one week after surgery, the mouse was habituated to head fixation in daily sessions of progressively increasing duration, up to 2 h. During head fixation, the mouse was positioned on a suspended bed and was periodically offered a drop of sweetened condensed milk as a reward. The habituated mouse readily consumed the reward and remained free to adjust its body position, occasionally displaying natural grooming behavior.

### Processing of dual-mode mouse cortex imaging

Raw dual-mode recordings consisted of alternating GCaMP fluorescence and 530 nm reflectance frames. The image sequence was separated into fluorescence and reflectance stacks according to the alternating acquisition order and saved as multi-page TIFF files for subsequent analysis.

Fluorescence (*F*) and reflectance (*R*) images were binned by 2 *×* 2 pixel averaging before processing. To compensate for slow channel-wide intensity drift, attributed primarily to LED heating, the fluorescence and reflectance channels were corrected independently using the entire recording. Let *X_c_*(*x, y, k*) denote the measured intensity at pixel (*x, y*) in channel frame *k*, where *c ∈ {F, R}* denotes fluorescence or reflectance and *N* is the total number of frames in that channel. For each pixel, the temporal-mean reference intensity was calculated over the full recording as

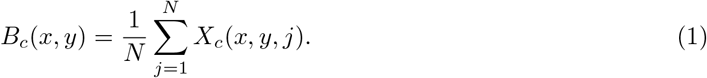

Time was expressed in channel frames and centered at the temporal midpoint of the recording:

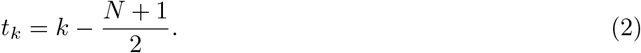

Thus, *t_k_* = 0 corresponds to the center of the recording. Under the linear-drift model described below, *B_c_*(*x, y*) serves as the pixel-specific reference intensity at this time point.

Pixels with *B_c_*(*x, y*) *≤* 1 in either channel were excluded to avoid normalization by near-zero reference intensities. The intersection of the fluorescence and reflectance validity masks was used to estimate the global drift in both channels. At each frame, the valid pixels were spatially averaged to obtain one global fluorescence trace and one global reflectance trace. No temporal smoothing was applied. For each channel, the global trace was fitted over the entire recording by least squares using

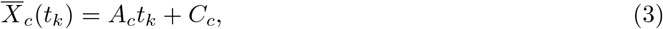

where *A_c_*is the fitted linear drift rate in intensity units per channel frame.

The fitted drift rate *A_c_*was assumed to be spatially uniform within each channel, whereas the reference intensity *B_c_*(*x, y*) was pixel dependent. The resulting time-dependent reference intensity was therefore defined as

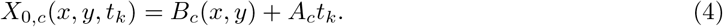

For the fluorescence and reflectance channels, respectively,

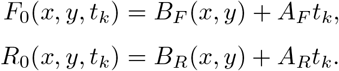

These functions represent the expected fluorescence and reflectance intensities after accounting for the fitted linear drift and provide the time-dependent reference intensities used for normalization at each pixel and time point.

The drift-normalized, hemodynamic-uncorrected fluorescence and reflectance changes were calculated as

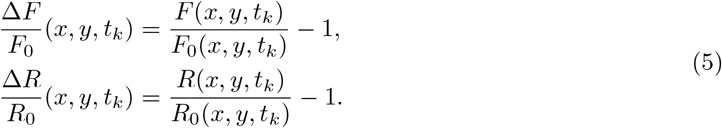

To reduce hemodynamic contamination of the GCaMP fluorescence signal, we applied a reflectance-based ratiometric correction adapted from wide-field optical mapping [19]. The 530 nm reflectance measurement provides a hemodynamic-sensitive signal dominated by changes in hemoglobin absorption and cerebral blood volume. Hemodynamic-corrected fluorescence was computed pixel-wise as

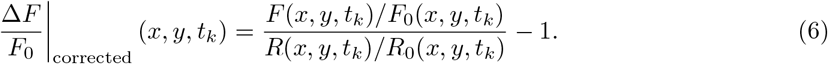

Here, *F* and *R* denote the frame-interleaved GCaMP fluorescence and 530 nm reflectance measurements, respectively. Each fluorescence frame was paired with the corresponding adjacent reflectance frame according to the acquisition order, without temporal interpolation. Pixels were retained only if their temporal-mean fluorescence and reflectance reference intensities exceeded 1 and their fitted time-dependent reference intensities remained greater than 10*^−^*^6^ throughout the recording. A numerical offset of 10*^−^*^6^ was added to denominators to prevent division by near-zero values. All other pixels were excluded from further analysis.

ROI time series were extracted from the drift-normalized fluorescence (Δ*F/F*_0_), reflectance (Δ*R/R*_0_), and hemodynamic-corrected fluorescence image stacks. Circular ROIs were manually defined on the average registered reference image and preferentially placed away from large surface vessels, where scattering and other non-absorptive optical effects may reduce the accuracy of the reflectance-based correction, which primarily compensates for absorption-related hemodynamic modulation [25]. For each frame, the ROI signal was computed as the mean value of all valid pixels within the ROI. The traces shown in Figure 6E were plotted directly from the normalized image stacks without additional baseline subtraction.

## Data Availability

Raw and processed experimental data, MATLAB processing code, Arduino trigger code, 3D-printable STL files, and supporting device files will be deposited in Zenodo at https://doi.org/10.5281/zenodo.21143140. The repository will include the scripts used for dual-channel stack splitting, mouse-cortex hemodynamic correction, and ROI trace extraction.

## Acknowledgments

We acknowledge funding from NIH (R01NS126596, U19NS123717, R01CA215059, R01NS140967) and an innovation grant from the Alex Lemonade Stand Foundation. Bradley C Rauscher was supported by a Ruth L. Kirschstein Predoctoral Fellowship F31NS145737. We acknowledge the support by the Neurophotonics Center at Boston University. We thank the Boston University Shared Computing Cluster for providing computational resources.

## Author Contributions

Q.D., G.H., and L.T. designed research; Q.D. and G.H. performed research; Q.D., B.C.R., M.V., B.W., and Z.C. contributed biological analysis; Q.D. and G.H. analyzed data; and Q.D. and L.T. wrote the paper.

## References

[1] J. Lichtman and JA. Conchello. Fluorescence microscopy. Nature Methods, 2:910–919, 2005.

[2] Natan T. Shaked, Stephen A. Boppart, Lihong V. Wang, and Jürgen Popp. Label-free biomedical optical imaging. Nature Photonics, 17:1031–1041, 2023.

[3] Kunal K. Ghosh, Laurie D. Burns, Eric D. Cocker, Axel Nimmerjahn, Yaniv Ziv, Abbas El Gamal, and Mark J. Schnitzer. Miniaturized integration of a fluorescence microscope. Nature Methods, 8(10):871–878, 2011.

[4] Daniel Aharoni, Baljit S. Khakh, Alcino J. Silva, and Peyman Golshani. All the light that we can see: a new era in miniaturized microscopy. Nature Methods, 16:11–13, 2019.

[5] Changliang Guo, Garrett J. Blair, Megha Sehgal, Federico N. Sangiuliano Jimka, Arash Bellafard, Alcino J. Silva, Peyman Golshani, Michele A. Basso, Hugh Tad Blair, and Daniel Aharoni. Miniscope-LFOV: A large-field-of-view, single-cell-resolution, miniature microscope for wired and wire-free imaging of neural dynamics in freely behaving animals. Science Advances, 9(16):eadg3918, April 2023.

[6] Pingping Zhao et al. Minixl: An open-source, large field-of-view epifluorescence miniscope enabling single-cell resolution and multi-region imaging in mice. Science Advances, 11:eads4995, 2025.

[7] Jia Hu, Arun Cherkkil, Daniel A. Surinach, Ibrahim Oladepo, Ridwan Hossain, Skylar Fausner, Kapil Saxena, Eunsong Ko, Ryan Peters, Michael Feldkamp, Pavan C. Konda, Vinayak Pathak, Roarke Horstmeyer, and Suhasa B. Kodandaramaiah. Pan-cortical cellular imaging in freely behaving mice using a miniaturized micro-camera array microscope (mini-mcam). Science Advances, 11(44):eadt3634, 2025.

[8] Yuanlong Zhang, Lekang Yuan, Qiyu Zhu, Jiamin Wu, Tobias Nöbauer, Rujin Zhang, Guihua Xiao, Mingrui Wang, Hao Xie, Zengcai Guo, Qionghai Dai, and Alipasha Vaziri. A miniaturized mesoscope for the large-scale single-neuron-resolved imaging of neuronal activity in freely behaving mice. Nature Biomedical Engineering, 8:754–774, 2024.

[9] Kyrollos Yanny, Nick Antipa, William Liberti, Sam Dehaeck, Kristina Monakhova, Fanglin Linda Liu, Konlin Shen, Ren Ng, and Laura Waller. Miniscope3D: optimized single-shot miniature 3d fluorescence microscopy. Light: Science & Applications, 9(1):171, 2020.

[10] Feng Tian et al. Deepinminiscope: Deep learning–powered physics-informed integrated miniscope. Science Advances, 11:eadr6687, 2025.

[11] Josefa R. Scherrer, Galen F. Lynch, Jie J. Zhang, and Michale S. Fee. An optical design enabling lightweight and large field-of-view head-mounted microscopes. Nature Methods, 20(4):546–549, 2023.

[12] Jimin Wu, Yuzhi Chen, Ashok Veeraraghavan, Eyal Seidemann, and Jacob T Robinson. Mesoscopic calcium imaging in a head-unrestrained male non-human primate using a lensless microscope. Nature Communications, 15(1):1271, 2024.

[13] Jesse K. Adams, Dong Yan, Jimin Wu, Vivek Boominathan, Sibo Gao, Alex V. Rodriguez, Soonyoung Kim, Jennifer Carns, Rebecca Richards-Kortum, Caleb Kemere, Ashok Veeraraghavan, and Jacob T. Robinson. In vivo lensless microscopy via a phase mask generating diffraction patterns with high-contrast contours. Nature Biomedical Engineering, 6(5):617–628, 2022.

[14] Yujia Xue, Ian G. Davison, David A. Boas, and Lei Tian. Single-shot 3D wide-field fluorescence imaging with a computational miniature mesoscope. Science Advances, 6(43):eabb7508, 2020.

[15] Yujia Xue, Qianwan Yang, Guorong Hu, Kehan Guo, and Lei Tian. Deep-learning-augmented computational miniature mesoscope. Optica, 9(9):1009–1021, 2022.

[16] Guorong Hu, Qilin Deng, Tianrui Qi, Zhixiong Chen, Bradley C. Rauscher, Nathan Chai, Daria Bogatova, Bethany Weinberg, Jesse Smith, Ian G. Davison, Martin Thunemann, Anna Devor, and Lei Tian. Bio-CM^2^: Distributed computational optics for cortex-wide cellular imaging. bioRxiv, 2026. Preprint. doi: 10.64898/2026.07.27.740823.

[17] Zhe Dong, Yu Feng, Keziah Diego, Austin M. Baggetta, Brian M. Sweis, Zachary T. Pennington, Sophia I. Lamsifer, Yosif Zaki, Federico Sangiuliano, Paul A. Philipsberg, Denisse Morales-Rodriguez, Daniel Kircher, Paul A. Slesinger, Tristan Shuman, Daniel Aharoni, and Denise J. Cai. Simultaneous two-color imaging with a dual-channel miniscope in freely behaving mice. Science Advances, 11(27):eadr6470, 2025.

[18] Jinyong Zhang, Feiyang Hong, Jiwon Kim, Konstantin Bakhurin, Namsoo Kim, and Henry H. Yin. A dual-color miniature endoscope for calcium imaging in behaving mice. iScience, 29(5):115514, 2026.

[19] Y. Ma et al. Wide-field optical mapping of neural activity and brain haemodynamics: considerations and novel approaches. Philosophical Transactions of the Royal Society B: Biological Sciences, 371:20150360, 2016.

[20] Matthieu P. Vanni and Timothy H. Murphy. Mesoscale transcranial spontaneous activity mapping in gcamp3 transgenic mice reveals extensive reciprocal connections between areas of somatomotor cortex. Journal of Neuroscience, 34(48):15931–15946, 2014.

[21] Anna Letizia Allegra Mascaro, Irene Costantini, Emilia Margoni, Giulio Iannello, Alessandro Bria, Leonardo Sacconi, and Francesco S. Pavone. Label-free near-infrared reflectance microscopy as a complimentary tool for two-photon fluorescence brain imaging. Biomedical Optics Express, 6:4483–4492, 2015.

[22] Mathew L. Rynes, Daniel A. Surinach, Samantha Linn, Michael Laroque, Vijay Rajendran, Judith Dominguez, Orestes Hadjistamoulou, Zahra S. Navabi, Leila Ghanbari, Gregory W. Johnson, Mojtaba Nazari, Majid H. Mohajerani, and Suhasa B. Kodandaramaiah. Miniaturized head-mounted microscope for whole-cortex mesoscale imaging in freely behaving mice. Nature Methods, 18(4):417–425, 2021.

[23] H. Hou, J. Wu, J. Liu, V. Boominathan, A. Shende, K. Goli, J. Carns, R.A. Schwarz, A.M. Gillenwater, P. Ramalingam, M.P. Salcedo, K.M. Schmeler, T.S. Tkaczyk, J.T. Robinson, A. Veeraraghavan, and R.R. Richards-Kortum. Deep-learning endomicroscope with large field-of-view and depth-of-field for real-time in vivo imaging of epithelial cancer hallmarks. Proceedings of the National Academy of Sciences of the United States of America, 123(20):e2602705123, 2026.

[24] Scott A. Prahl. Optical absorption of hemoglobin. Oregon Medical Laser Center, 1999. Available at: https://omlc.ogi.edu/spectra/hemoglobin/.

[25] Patrick R. Doran, Natalie Fomin-Thunemann, Rockwell P. Tang, Dora Balog, Bernhard Zimmerman, Kıvılcim Kılıç, Emily A. Martin, Sreekanth Kura, Harrison P. Fisher, Grace Chabbott, Joel Herbert, Bradley C. Rauscher, John X. Jiang, Sava Sakadzic, David A. Boas, Anna Devor, Ichun Anderson Chen, and Martin Thunemann. Widefield in vivo imaging system with two fluorescence and two reflectance channels, a single scmos detector, and shielded illumination. Neurophotonics, 11(3):034310, 2024.

[26] James F. Morley, Heather R. Brignull, Jill J. Weyers, and Richard I. Morimoto. The threshold for polyglutamine-expansion protein aggregation and cellular toxicity is dynamic and influenced by aging in Caenorhabditis elegans. Proceedings of the National Academy of Sciences, 99(16):10417– 10422, 2002.

[27] Bradley C Rauscher, Natalie Fomin-Thunemann, Sreekanth Kura, Patrick R Doran, Pablo D Perez, Kıvılcım Kılıç, Emily A Martin, Dora Balog, Nathan X Chai, Francesca A Froio, et al. The neurovascular impulse response function differentially reflects intrinsic neuromodulation across cortical regions. Nature neuroscience, pages 1–9, 2026.

[28] Ken Y. Chan, Min J. Jang, Bryan B. Yoo, Alon Greenbaum, Namita Ravi, Wei-Li Wu, Luis Sánchez-Guardado, Carlos Lois, Sarkis K. Mazmanian, Benjamin E. Deverman, and Viviana Gradinaru. Engineered AAVs for efficient noninvasive gene delivery to the central and peripheral nervous systems. Nature Neuroscience, 20(8):1172–1179, 2017.

[29] Glenn J. Goldey, Demetris K. Roumis, Lindsey L. Glickfeld, Aaron M. Kerlin, R. Clay Reid, Vincent Bonin, Dorothy P. Schafer, and Mark L. Andermann. Removable cranial windows for long-term imaging in awake mice. Nature Protocols, 9(11):2515–2538, 2014.

[30] Kıvılcım Kılıç, Michèle Desjardins, Jianbo Tang, Martin Thunemann, Smrithi Sunil, Şefik Evren Erdener, Dmitry D. Postnov, David A. Boas, and Anna Devor. Chronic cranial windows for long term multimodal neurovascular imaging in mice. Frontiers in Physiology, 11:612678, 2021.

